# The generation of the first chromosome-level de-novo genome assembly and the development and validation of a 50K SNP array for North American Atlantic salmon

**DOI:** 10.1101/2022.09.28.509896

**Authors:** Guangtu Gao, Geoffrey C. Waldbieser, Ramey C. Youngblood, Dongyan Zhao, Michael R. Pietrak, Melissa S. Allen, Jason A. Stannard, John T. Buchanan, Roseanna L. Long, Melissa Milligan, Gary Burr, Katherine Mejía-Guerra, Moira J. Sheehan, Brian E. Scheffler, Caird E. Rexroad, Brian C. Peterson, Yniv Palti

## Abstract

Given the genetic and genomic differences between Atlantic salmon of European origin and North American (N.A.) origin, it is crucial to develop unique genomic resources for each lineage. Here we describe the resources that we recently developed for genomic and genetic research in N.A. Atlantic salmon. Firstly, a new single nucleotide polymorphism (SNP) database for N.A. Atlantic salmon consisting of 3.1 million putative SNPs was generated using data from whole genome resequencing of 80 N.A. Atlantic salmon individuals; Secondly, a high density 50K SNP array enriched for the genic regions of the genome and containing three sex determination and 61 continent of origin markers was developed and validated; Thirdly, a genetic map composed of 27 linkage groups with 36K SNP markers, was generated from 2,512 individuals in 141 full-sib families; Finally, a chromosome level de-novo assembly of a male N.A. Atlantic salmon genome was generated using PacBio long-reads. Information from Hi-C proximity ligation sequences and Bionano optical mapping was used to concatenate the contigs into scaffolds. The assembly contains 1,755 scaffolds and only 1,253 gaps, with a total length of 2.83 Gb and N50 of 17.2 Mb. A BUSCO analysis detected 96.2% of conserved Actinopterygii genes in the assembly and the genetic linkage information was used to guide the formation of 27 chromosome sequences. In contrast, the karyotype of the European Atlantic salmon lineage is composed of 29 chromosomes. Comparative analysis with the reference genome assembly of the European Atlantic salmon confirmed that the karyotype differences between the two linages are caused by a fission in chromosome Ssa01 and three chromosome fusions including the p arm of chromosome Ssa01 with Ssa23, Ssa08 with Ssa29 and Ssa26 with Ssa28. The genomic resources we have generated for Atlantic salmon provide a crucial boost for genetic research and for management of farmed and wild populations in this highly valued species.

## Introduction

Atlantic salmon (*Salmo salar*) in Northeastern US and Eastern Canada has high economic value for the sport fishing and aquaculture industries and is of significant ecological and social impact (Council 2004). Atlantic salmon of North American (N.A). origin is also being farmed in Tasmania, Australia (Kijas et al. 2017) and hence also an important commodity in that continent. The historic declines of wild Atlantic salmon populations in the state of Maine, USA, led to the listing of wild or natural Atlantic salmon from eight rivers as endangered under the federal Endangered Species Act (Council 2004). Evidence from microsatellite DNA markers has clearly demonstrated that Atlantic salmon of N.A. origin is genetically distinct from European Atlantic salmon (King et al. 2001; King et al. 2005; Council 2002). This led to a judicial ruling that strictly regulated farming of Atlantic salmon in Maine and eastern Canada, banned the farming of European and genetically modified salmon strains in these areas, and required routine genetic certification of the North American aquaculture stocks (Council 2004). More recent studies that used whole genome scans with high density SNP assays provided further evidence for the genomic differentiation between the North American and European lineages (Lehnert et al. 2020).

The genome size estimate for Atlantic salmon based on cellular DNA content is 3.0 Gbp (Hardie and Hebert 2003). Although it is similar in size to that of most mammals, its architecture and organization are more complex. All ray-finned fish share an additional (3R) round of ancestral genome duplication not found in mammals and birds, but the salmonids have undergone an additional, more recent (4R) whole genome duplication event (Allendorf and Thorgaard 1984; Lien et al. 2016). Approximately 25% of the European Atlantic salmon genome assembly in seven pairs of chromosome arms with elevated sequence similarity (>90%) and patterns of tetrasomic inheritance, and in total, 94.4% of the genome is composed of 98 pairs of collinear blocks (196 regions) of high sequence similarity known as homeologous chromosome regions (Lien et al. 2016). In addition, nearly 60% of the Atlantic salmon genome contains repetitive sequences (Lien et al. 2016).

Single-nucleotide polymorphisms (SNP) are highly abundant markers broadly distributed in animal genomes. High density SNP arrays are used for collecting large amount of genome-wide genotype data. This information is useful for dissecting the genetic basis of quantitative traits in agriculture and for implementing models of genomic selection, which has revolutionized the field of selective breeding over the past decade (Meuwissen et al. 2016). High density SNP arrays are publicly available for Atlantic salmon of European origin (Houston et al. 2014). The N.A. Atlantic salmon is a different sub-species with substantial genomic differences from the European sub-species. The haploid chromosome number of N.A. Atlantic salmon is N=27 compared to N=29 for European salmon (Lubieniecki et al. 2010; Brenna-Hansen et al. 2012), and only half of the SNPs from the arrays designed for the European salmon are informative for N.A. salmon (John Buchanan, unpublished data) (Yáñez et al. 2016). Other researchers and commercial breeding companies have developed optimized SNP chips for salmon of N.A. origin based on experimental information from the larger European salmon arrays, but those are currently not available for the public due to intellectual property constrains from competitive commercial interests (John Buchanan, unpublished data) (Kijas et al. 2018).

Recently we initiated efforts to generate genomic resources for the breeding program at the USDA-ARS National Cold Water Marine Aquaculture Center (NCWMAC) in Franklin, Maine. We used high coverage whole genome Illumina resequencing for SNP discovery in 80 N.A. Atlantic salmon individuals from three aquaculture stocks that are propagated in the NCWMAC (Gao et al. 2020). Overall, we discovered 6.6 million SNP markers, including more than 1.5 million markers with minor allele frequency (MAF) ≥ 0.25. In addition, we identified 5,822 candidate markers that can potentially be used to distinguish between N.A. and European Atlantic salmon by comparing genotypes of the 80 N.A. Atlantic salmon with publicly available whole genome sequence information from 31 Atlantic salmon representing a diversity of European populations. Although this SNP database provided a foundational resource for designing new SNP genotyping arrays specifically tailored for genomic and genetic analyses in N.A. Atlantic salmon, we also found a high percentage of sequence mismatch from the alignment of N.A. salmon sequencing reads to the European salmon reference genome assembly. This indicated that a reference genome from N.A. salmon can improve the quality and genome coverage of SNP discovery in these fish.

We report here the first *de-novo* genome assembly for the North American Atlantic salmon sub-species, generated from a male fish from the St. John River aquaculture strain. Advances in long-read DNA sequencing technologies, together with the development of new assembly algorithms and pipelines, have recently enabled improved genome assemblies for complex salmonid genomes like the rainbow trout (Gao et al. 2021).The new genome assembly for Atlantic salmon was generated using PacBio long-read sequencing technology followed by scaffolding with Hi-C contact maps and Bionano optical mapping. The result was high quality, contiguous chromosome sequences. The contigs of the new assembly were used as reference to generate a new SNP discovery dataset for N.A. Atlantic salmon from which SNPs were selected for a 50K Axion SNP array, including markers that can be used for continent of origin classification (Europe or North America) and for sex determination by testing for presence of the sdY gene in the genome in the individual fish. The new SNP array was validated by genotyping pedigreed fish from the USDA breeding program and generating a linkage map that facilitated anchoring and ordering of the sequence scaffolds in 27 chromosome sequences. The reference genome and high-density SNP array provide very important resources for more precise breeding in aquaculture, and for comparative genomics research on the ecology and evolution of Atlantic salmon.

## Materials and Methods

### SNP discovery

Genomic DNA was extracted from the fin clips of 80 N.A. Atlantic salmon selected from three aquaculture stocks that are currently propagated at the USDA-ARS National Cold Water Marine Aquaculture Center (NCWMAC) in Franklin, Maine. These stocks represent three distinctive strains of N.A. Atlantic salmon - the Gaspe of New Brunswick (GNB) is a landlocked strain and the Penobscot River (PR) strain was originated from an endangered N.A. Atlantic salmon population. The St. John River (SJR) strain is of great economic importance as the origin of commercial strains in the Northeast US and Canadian aquaculture industry, and it is also the primary species used in the selective breeding program in NCWMAC. Genomic DNA was extracted from 80 fish - 16 GNB, 53 SJR, and 11 PR strain. Whole genome shotgun sequencing was performed using Illumina NovaSeq (paired-end 2 x 150 nucleotides) as previously described (Gao et al. 2020). All raw sequence data can be found in the NCBI SRA repository (BioProject accession PRJNA559280).

Quality of raw sequencing reads was assessed using FASTQC. Adapter sequences and low-quality bases were removed using Trimmomatic (LEADING:10 TRAILING:10 SLIDINGWINDOW:4:15 MINLEN:30) (Bolger et al. 2014) and only surviving reads in pairs were used for downstream SNP discovery. Cleaned reads were aligned to an earlier draft of the Canu contigs assembly of the North American Atlantic salmon genome assembly (GenBank Accession GCA_021399835.1, reported here for the first time) using BWA mem with the default parameters (Li 2013). Duplicated alignments were marked using PicardTools v2.19.2 (http://broadinstitute.github.io/picard). Reads in regions of detected indels were realigned to remove alignment artifacts and base quality scores were recalibrated based on the realignments to generate analysis-ready reads in bam format using GATK v3.8.1 (Poplin et al. 2018). Raw variants, including SNPs and indels, were generated using HaplotypeCaller and GenotypeGVCF commands in GATK v3.8.1 (Poplin et al. 2018). SNPs were retained if they met the following filtering criteria: 1) located more than 5 bp from an indel; 2) QUAL > 30; 3) minimum and maximum read depths of 100 and 3000, respectively; 4) for each sample, at least one read supporting reference allele and two reads supporting the alternative allele; 5) no missing genotype per SNP position; and 6) MAF > 0.3.

### Selection of SNPs for the 50K Array

The annotation of the European Atlantic salmon reference genome was used as a proxy to locate genic regions in the North American salmon genome. Whole genome alignments between sequence contigs from the North American and European Atlantic salmon genome were generated using minimap2 (Li 2018). Alignments passing the following requirements were retained for downstream analyses: 1) alignments to the 29 chromosomes of the European salmon genome and contigs of the North American salmon genome assembly that are longer than 5,000 bp; 2) minimum alignment length of 5,000 bp; 3) minimum mapping quality of 40; 4) a North American salmon contig has to have the “assigned” European salmon chromosome (using the sum of base pairs as a voting system, discard ties); 5) only one North American salmon contig was kept if multiple contigs aligned to the same region of the European genome. To select 100K SNPs for technical ranking by the SNP array manufacturer, ~35 SNPs were selected in each 1 Mb window, with a minimum distance between adjacent SNPs of 1.5 Kb and prioritized SNPs in genic regions based on the alignments to the European genome annotation. The dataset of 100K SNPs included 8,654 from one of the European-based high density SNP arrays (Houston et al. 2014) that were found to be high quality and polymorphic in the North American stock of the Mowi ASA aquaculture company. The SNP information for those markers was provided courtesy of Serap Gonen and Matthew Baranski and the original Affymetrix SNP IDs for those markers were retained in the new 50K SNP array.

The ~100K SNPs were then evaluated by the array manufacturer (ThermoFisher, USA) for best compatibility with the SNP array platform, from which 49,804 were selected as the core SNPs of the 50K Axiom SNP array. The total number of markers on the array is 49,880 which also included 72 SNPs that can potentially be used for continent of origin (COO) classification, including 44 nuclear markers and 20 mitochondrial markers identified previously (Gao et al. 2020) and eight nuclear markers that were previously found to be informative for COO classification by the Center for Aquaculture Technologies (unpublished data). Four additional sequence probes from the sdY gene were also added to the array for sex determination analysis. Those probes were previously included and tested in other Axiom SNP arrays for Atlantic salmon (Kijas et al. 2018). Presence of the sequences compatible with the sdY probes would indicate that the fish is a male and absence that it is a female.

### Genotyping and validation of the SNP array

Fin tissues were collected, and genomic DNA was extracted from 2,512 pedigreed fish from the SJR strain of the USDA breeding program. Additional DNA samples (Table S1) were used to evaluate the continent of origin markers in the Axiom array. Genomic DNA samples were fluorescently labeled, hybridized to the SNP array and scanned for initial genotype calls (Neogen, Lincoln, NE) according to the Axiom genotyping procedures described by Affymetrix. For final genotyping calls and quality control analyses we used the Affymetrix Power Tools (APT) and SNPolisher software packages following the procedures and practices suggested by the Affymetrix Axiom array genotyping manual and in the same way we previously described for the rainbow trout 57K Axiom array (Palti et al. 2015).

### Linkage Map

The fish used in the linkage analysis were from the St. John River stock of N. A. Atlantic salmon (NCWMAC-USDA). In total we genotyped 2,512 fish from 141 full-sib families, including 230 parents and 2,282 offspring. The program Lep-map3 (Rastas 2017) was used for generating the linkage map with the same iterative step-wise approach used for the rainbow trout linkage map (Gonzalez-Pena et al. 2016). Briefly, the markers were separated into 27 linkage groups by running the SeparateChromosomes2 module at LOD score 110. More markers were added to the linkage groups by running the JoinSingles2All module with a more relaxed LOD threshold. Finally, markers were ordered in each linkage group by running the OrderMarkers2 module. Linkage groups were aligned with the karyotype chromosomes by co-mapping the flanking sequences of SNP markers from this linkage map and the map of Brenna-Hansen et al. (Brenna-Hansen et al. 2012) to the 27 primary sequence scaffolds of the new N.A. Atlantic salmon genome assembly. Two programs were used to map the flanking sequences of the SNP markers to the genome assembly sequence scaffolds. The program NovoAlign (http://www.novocraft.com/products/novoalign/) with the parameter -t 60 was used for the SNP markers of this linkage map, and Blastn (Zhang et al. 2000) with the parameters of, -evalue 1.0e-5, -perc_identity 80, and -qcov_hsp_perc 80, was used for the markers of Brenna-Hansen et al. (2012).

### Genome assembly - DNA extraction and sequencing

DNA from a single male (sample NCWMAC SJR YC14-15 Fam3) from the USDA St. John River stock of N. A. Atlantic salmon was extracted using the Qiagen DNeasy kit from a fresh blood sample collected in tri-potassium EDTA. The blood was stored on ice until DNA extraction within 24 hours to enhance the yield of high molecular weight DNA for long-read sequencing. The DNA sample was sequenced at the core facility of the University of Delaware (Newark, Delaware, USA) using 29 SMRT cells of the PacBio RS-II system in continuous long reads (CLR) mode, which generated approximately 312 Gb of sequence data (104x coverage). The PacBio read length N50 was ~33 kb and the average read length was ~19 kb. For polishing the genome assembly, ~2.1 billion paired-end reads (2 x150bp) of Ilumina short-reads were generated by Admera Health (South Plainfield, NJ, USA) from the same DNA sample. The short-read sequences were processed with the program cutadapt to remove the Illumina sequencing adaptors producing a total of ~316 Gb (~105x genome coverage). In general, high genome coverage above 30x is desired for polishing with Illlumina reads.

### Construction of the assembly contigs

The initial assembly of sequence contigs was generated from the PacBio long-reads using the Canu (Version 1.8) assembler (Koren et al. 2017) with the options of correctedErrorRate=0.085, corMhapSensitivity=normal, and ovlMerDistinct=0.975. After the “correction” and “trimming” steps, which involve alignments of reads, error correction, trimming of poor sequence quality segments and removal of overlapping reads after those corrections, Canu retained 2.64 million high quality reads with 103.5 Gb (~34.5x) total length from the raw PacBio reads. The trimmed reads were then assembled into 14,845 contigs.

Next, the programs purge_haplotigs (Roach et al. 2018) and purge_dups (Guan et al. 2020) were used to correct the Canu assembly errors and identify regional duplications due to high heterogeneity in some of the genomic regions. First, we ran purge_haplotigs to identify artefactual contigs caused by very low or very high coverage detected by mapping of the original PacBio long reads to the canu contigs. As a result, 1,394 canu contigs were characterized as artefacts, and thus removed from further analysis. Second, the retained sequences were processed with purge_dups to identify highly overlapping haplotigs. With this step, 2,263 and 11,977 sequences were identified as the primary contigs and haplotigs, respectively. This information was then used in construction of the genome assembly scaffolds.

### Construction of the assembly scaffolds and chromosome sequences

To improve the contiguity of the assembly, we first used Bionano optical mapping with the Saphyr platform (Bionano Genomics, San Diego, CA, USA) to scaffold the assembled contigs. High molecular weight DNA was extracted in agarose gel plugs from fresh blood of the same Atlantic salmon fish by Amplicon Express (Pullman, Washington) and the extracted DNA molecules were labeled with the Direct Label and Stain (DLS) DNA Labeling kit utilizing the Direct Label Enzyme (DLE-1) (Bionano Genomics, San Diego, CA, USA). A total of 8.5 million DNA molecules were generated with a total length of 1.2 Tb (average length 141 Kb). The raw mapping data were processed with Bionano Solve software (Solve3.3_10252018) (https://bionanogenomics.com/support-page/bionano-access-software/) to generate a de novo Bionano assembly guided by the contigs of the PacBio sequences assembly. About 1.9 million molecules with N50 of ~305 Kb (~150x genome coverage) and average label density of 10 sites per 100 Kb were selected from the raw data after the filtering step, and 1,881 Bionano genome maps were generated with a total length of ~3.09 Gb and N50 of ~5.1 Mb. The Bionano genome maps and the 14,240 primary contigs and haplotigs were then scaffolded using the Bionano hybrid scaffold pipeline. This resulted in the joining of 2,174 contigs (~2.61 Gb) into 817 scaffolds (~2.65 Gb) and breaking of 140 contigs that were considered by the Bionano pipeline to be chimeric joins. The BioNano software utilized 13 bp gaps to indicate the presence of small gaps of undetermined size or undetermined flanking sequence overlaps (Gao et al. 2021), and the sequences flanking these gaps were manually corrected. This resulted in the removal of 21 sequence overlaps and the total length of the scaffolds was reduced to ~2.63 Gb with N50 of ~14.2 Mb. Finally, 1,195 primary contigs (~0.2 Gb) that were not joined with other sequences and not > 3,000 bp were added to the BioNano scaffolds for further construction of the assembly. Altogether, the BioNano assembly was composed of 2,012 sequence scaffolds with total length of ~2.84 Gb and N50 of ~11.94 Mb.

To improve the accuracy of the BioNano scaffolds, we first used gcpp (http://github.com/PacificBiosciences/GenomicConsensus) to polish the sequences. The PacBio raw reads were first mapped to the BioNano scaffolds using minimap2 (Version 2.15-r915) (Li 2018), and the scaffolds were polished using the “arrow” algorithm in gcpp based on the alignment results. To further polish the scaffolds, the Illumina short-read sequences were aligned to the scaffolds using BWA-MEM (Version 0.7.17-r188) (Li 2013) with the default parameters and the genomic variants were called using the freebayes (Version 1.3.1) program (Garrison and Marth 2012) with the default parameters. Variants with quality score > 1 and the alternative allele in the Illumina sequence reads were identified and the scaffolds were corrected based on the selected variants using bcftools (Li et al. 2009). This polishing procedure was performed twice to increase the accuracy of the genome assembly.

The Hi-C proximity ligation sequence data was used to generate chromosome scale scaffolds from the polished Bionano scaffold assembly. Fresh blood (100 ul) was collected in ethanol (750 ul) and immediately placed in dry ice and shipped. The Hi-C library was prepared with the Arima HiC kit (April 2018) using restriction enzymes that cut at GATC and GANTC recognition sites and sequenced to produce 833 million Illumina paired-end reads (2×150 bp) (Arima Genomics, San Diego, CA, USA). The reads were aligned to the Bionano scaffolds and the alignments were filtered following the Arima-HiC mapping pipeline (http://github.com/ArimaGenomics/mapping_pipeline). Briefly, the forward and reverse sequence reads of the Illumina pair-ends were mapped separately to the Bionano hybrid scaffolds using the BWA-MEM aligner (Li 2013). The chimeric 3’ side of the aligned reads that crossed the ligation junctions was removed and the reads that were mapped from both ends with the mapping quality score higher than 10 were retained. PCR duplicates in the Hi-C sequences were identified and removed using the Picard Toolkit (http://broadinstitute.github.io/picard/). After these filtering steps, ~300 million paired read alignments were retained, including ~79 million inter-scaffold alignments. The paired-read alignment data were then processed with SALSA (Ghurye et al. 2019) which broke 251 chimeric scaffolds and merged the polished Bionano assembly scaffolds into 1,663 Hi-C scaffolds with total length of ~2.83 Gb and N50 of ~37.9 Mb.

Linkage information from the new genetic maps was used to generate chromosome sequences from the Hi-C assembly scaffolds. The program NovoAlign (http://www.novocraft.com/products/novoalign/) was used to map the flanking sequences of each SNP marker to the scaffolds and the genetic linkage information was used to anchor the scaffolds to chromosomes and then to order, orient and concatenate the scaffolds into 27 chromosomes. With the guidance of the linkage information, 56 Hi-C scaffolds were broken into 150 smaller scaffolds at 94 Hi-C join sites due to incorrect ordering or wrong orientation of the joined scaffolds, or due to the insertion of a smaller scaffold at the join site of the larger scaffold.

### Alignment with the assembly of the European salmon genome

To align the chromosome sequences from the new assembly (GCA_021399835.1) to the chromosome sequences of the European Atlantic salmon genome assembly, repeat masked chromosome sequences of assembly Ssal_v3.1 (GCA_905237065.2) were used as the reference, and Blastn was used as the aligner with the parameters -evalue 1.0e-100, -word_size 51. All alignments longer than 2,000 bp were selected and plotted with the plot function of the program R.

## Results and Discussion

### New SNP database for N.A. Atlantic salmon

A new SNP database of ~3.1 million SNPs mapped to contigs from an earlier draft of the new N.A. Atlantic salmon genome assembly (GCA_021399835.1) was generated using GATK v3.8.1 (Poplin et al. 2018) as described in the methods. Of those SNPs, ~2.3M can be uniquely mapped to the chromosome sequences in the current assembly. The new database of ~2.3M SNPs uniquely mapped to a single chromosome position is available for download including the chromosome position and alternate alleles for each SNP (**Supplemental File S1**). To generate this file, we mapped the SNPs in retrospect to the final version of the chromosome sequences. We used the Novoalign program (www.Novocraft.com) to align 30bp flanking sequence from each side of the SNP to find exact matches to the genome assembly chromosome sequences. Of the 0.8M SNPs that were not uniquely mapped, 0.2M mapped to unplaced scaffolds, 0.4M were mapped to multiple locations in the chromosomes (MSVs), and 0.2M were duplicated (PSVs) as each allele mapped uniquely to a different position in the chromosome sequences.

### Genotyping and validation with the 50K SNP array

The SNPs were validated with the pedigreed population used to produce the linkage map (N=2,512). The marker categories were based on the Affymetrix Axiom array cluster properties. The total number of putative SNPs placed on the chip was 49,880. A total of 36,206 SNPs (73%) were categorized as high quality and polymorphic and an additional 1,825 SNPs (4%) as high quality but monomorphic, using the default quality filtering of the Affymetrix SNPolisher software. The number of SNPs passing the quality filtering steps is summarized in **Table 1.** Among the polymorphic high-resolution markers, average and median MAF were 0.32 and 0.34, respectively, and average and median heterozygosity were 40% and 45%, respectively. The markers distribution by MAF is shown in **Figure 1**.

**Figure 1.**
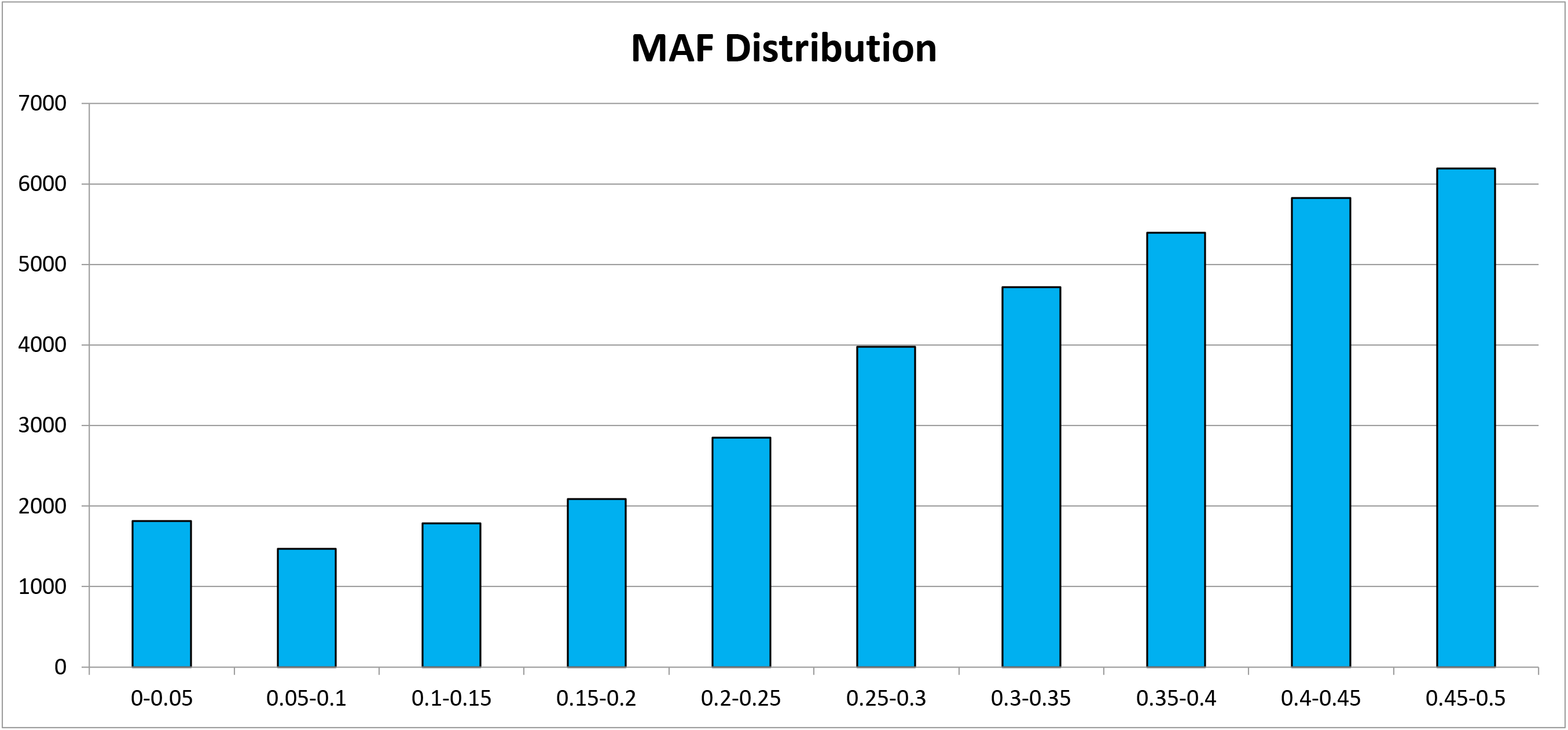
Polymorphism of SNPs from the Axiom 50K array. Minor allele frequency (MAF) distribution among the ~36K of polymorphic high-resolution SNPs

**Table 1.** Validation of the SNPs with DNA samples from North American Atlantic salmon (N=2,512). The marker categories are based on the Affymetrix Axiom array cluster properties.

| Category | SNP Markers |
| --- | --- |
| **Total SNPs on the Array** | 49,880 (100%) |
| **Call Rate <97%** | 5,421 (10.9%) |
| **No Minor Homozygotes** | 797 ( 1.6%) |
| **Off Target Variant** | 25 (0.05%) |
| **Other** | 5,586 (11.2%) |
| **Monomorphic High Resolution** | 1,825 ( 3.7%) |
| **Hemizygous (Mitochondrial)** | 20 (0.04%) |
| **Polymorphic High Resolution** | **36,206 (72.6%)** |

### Continent of origin markers

Forty one SNPs from the nuclear chromosomes and 20 mitochondrial SNPs were confirmed to be useful for identification of continent of origin (COO) using 60 samples from European origin populations and 22 from N.A. origin. These same samples were part of the panel that was originally used to test the King Seven microsatellites (King et al. 2005; King et al. 2001). Also, the 2,512 SJR aquaculture strain fish of N.A. origin that were used for general validation of the SNP array quality and for generating the linkage map were used to validate COO markers. The populations sampled are listed in **Table S1** and the markers validated for COO identification including the allele associated with each COO and their genome positions are listed in **Table S2**. The nuclear COO markers were not randomly distributed in chromosomes. The flanking sequences of 39 of the 41 COO nuclear markers were mapped to 13 of the 27 chromosomes with 7 and 6 markers mapped to chromosomes 12 and 14, respectively (Table S2). This nonrandom distribution may indicate association of genes on those two chromosomes with the genetic differentiation between the two lineages of Atlantic salmon. However, in contrast to our findings of enriched presence of COO-associated SNP markers on chromosomes 12 and 14, Lehnert et al. (Lehnert et al. 2020) identified genomic regions associated with high differentiation between the two lineages on chromosomes 06, 13, 16 and 19.

One of the N.A. origin fish from Gander – Jonathan’s Brook (sample NF2-015) had European mitochondria genotypes and N.A. nuclear genotypes, indicating possible introgression of the European mitochondrial genome due to ancestral hybridization. Evidence for introgression of European genotypes into North American populations was previously reported for Atlantic salmon in Newfoundland (Bradbury et al. 2015). Also, European mtDNA alleles have been confirmed in several Newfoundland populations of Atlantic salmon (Ian Bradbury, personal communication).

### Sex determination markers

Three markers derived from Atlantic salmon sdY sequences (Kijas et al., 2018) were placed on the array in duplicates (i.e. two Probset IDs for each SNP ID). The positions of the flanking sequences in the sdY gene sequence, Probset IDs and the flanking sequences are listed in **Table S3**. The alternative alleles for SNP ID Affx-158802265, Affx-158802258, and Affx-158802275 were C/T, C/T and G/T, respectively. Homozygous T alleles correlated with the absence of the sdY sequence (female) and homozygous alternative alleles or heterozygous genotypes correlated with sdY presence (male). To evaluate the accuracy of the sex determination markers we genotyped 135 female and 95 male parents of the families used for the linkage analysis and found 100% concordance of sex phenotype with the sdY marker genotypes.

### Linkage Map

A total of 36,158 polymorphic high-resolution markers were mapped to 27 linkage groups with an average of 1,339 markers per group (**Supplemental File S2**). The number of markers per linkage group ranged from 712 to 2,224. The distribution of markers in linkage groups labeled to match the nomenclature of the European Atlantic salmon karyotype is shown in **Figure 2**. Similar to previous reports for Atlantic salmon and rainbow trout (Pearse et al. 2019; Lien et al. 2016), the pattern of genetic distance in the paternal map was composed of high levels of recombination in the telomeric regions and high interference in the centromeric region and throughout most of the chromosome. Conversely, the typical maternal pattern of recombination in acrocentric chromosomes demonstrated higher rates of recombination near the centromere and then a steady level of recombination throughout the rest of the chromosome that gradually decreased to a plateau in the telomere region. In the metacentric chromosome the maternal pattern demonstrated strong recombination interference in the centromeric region and then a similar pattern to the acrocentric chromosome in the direction of either telomere whereas the paternal chromosome was demonstrated to have high levels of interference except near both telomeres (**Figure 3)**.

**Figure 2.**
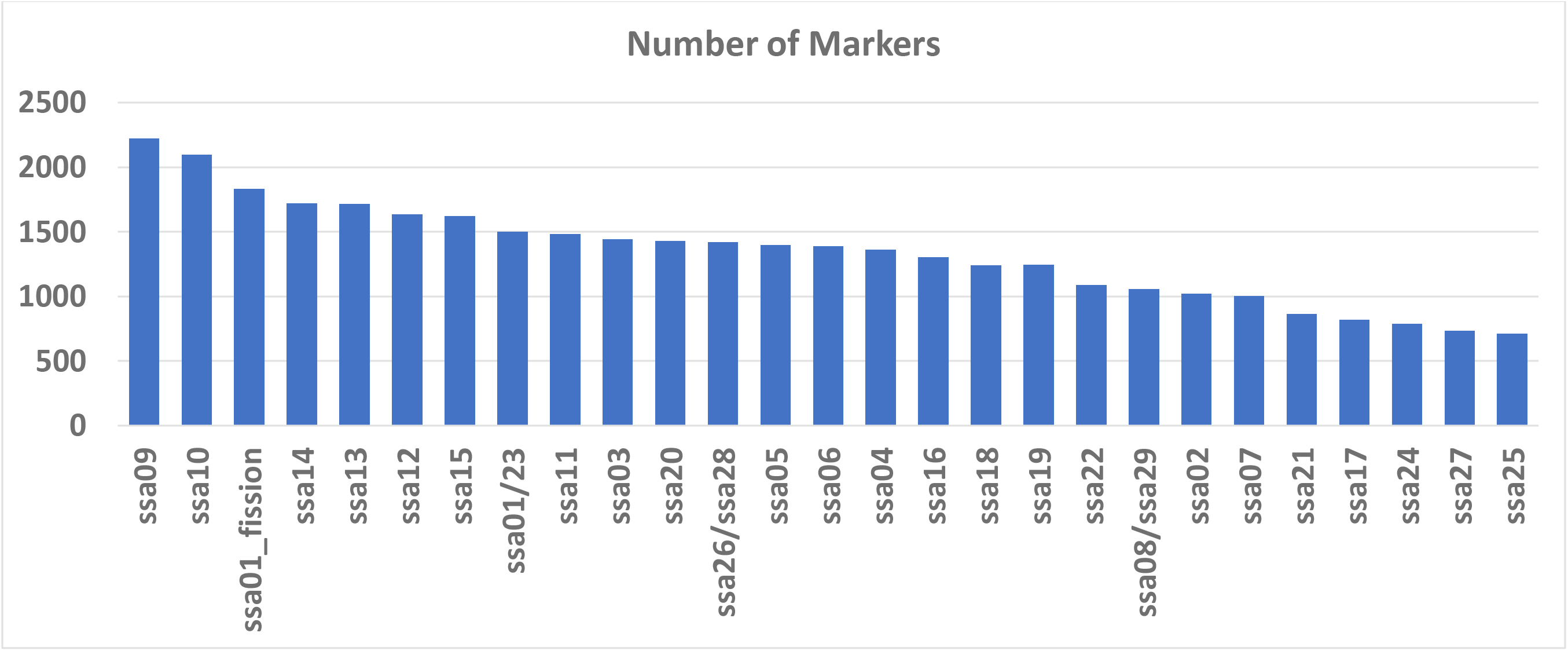
Distribution of the linkage map markers in chromosomes. The chromosome axis is in the order of linkage groups from LG1 to LG27. Chromosomes are labeled to match the nomenclature of the European Atlantic salmon karyotype.

**Figure 3.**
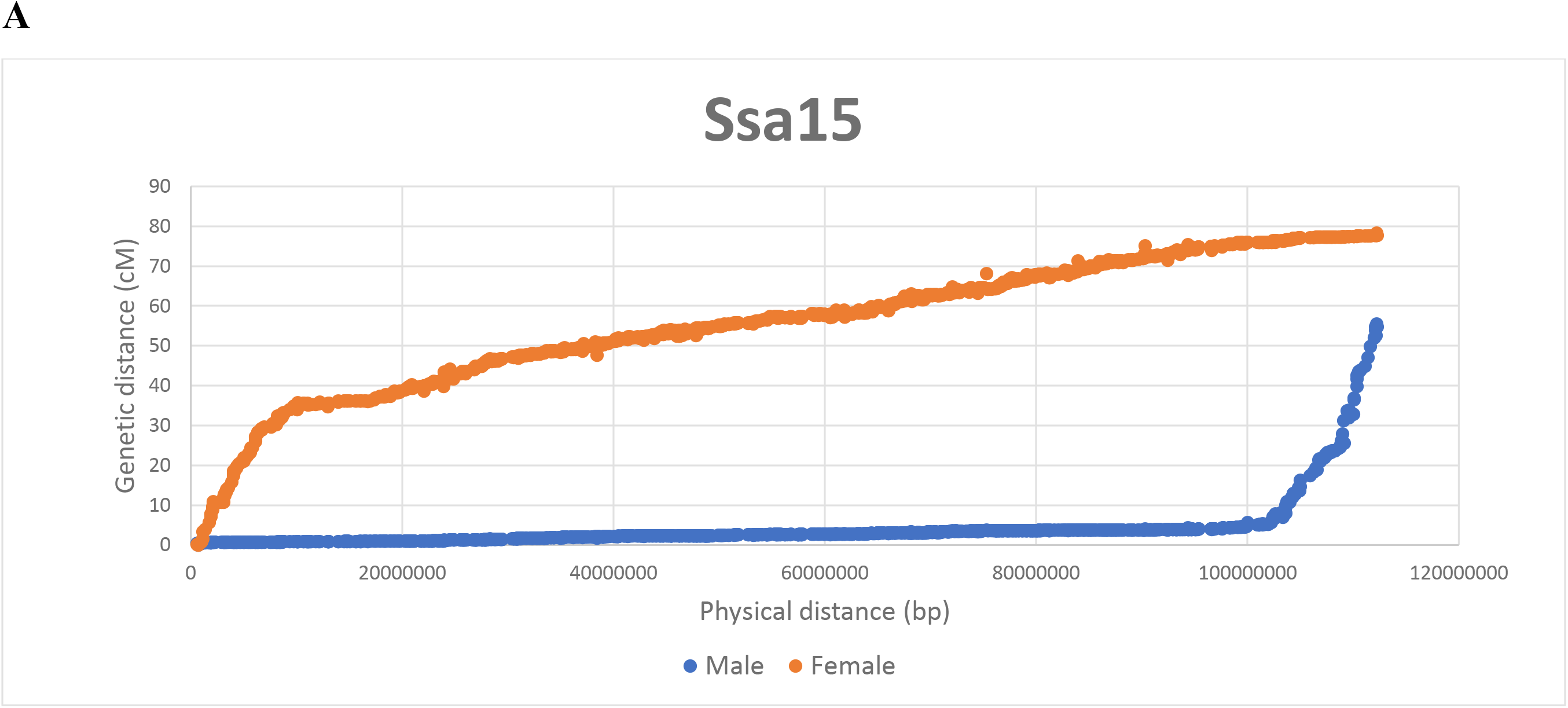
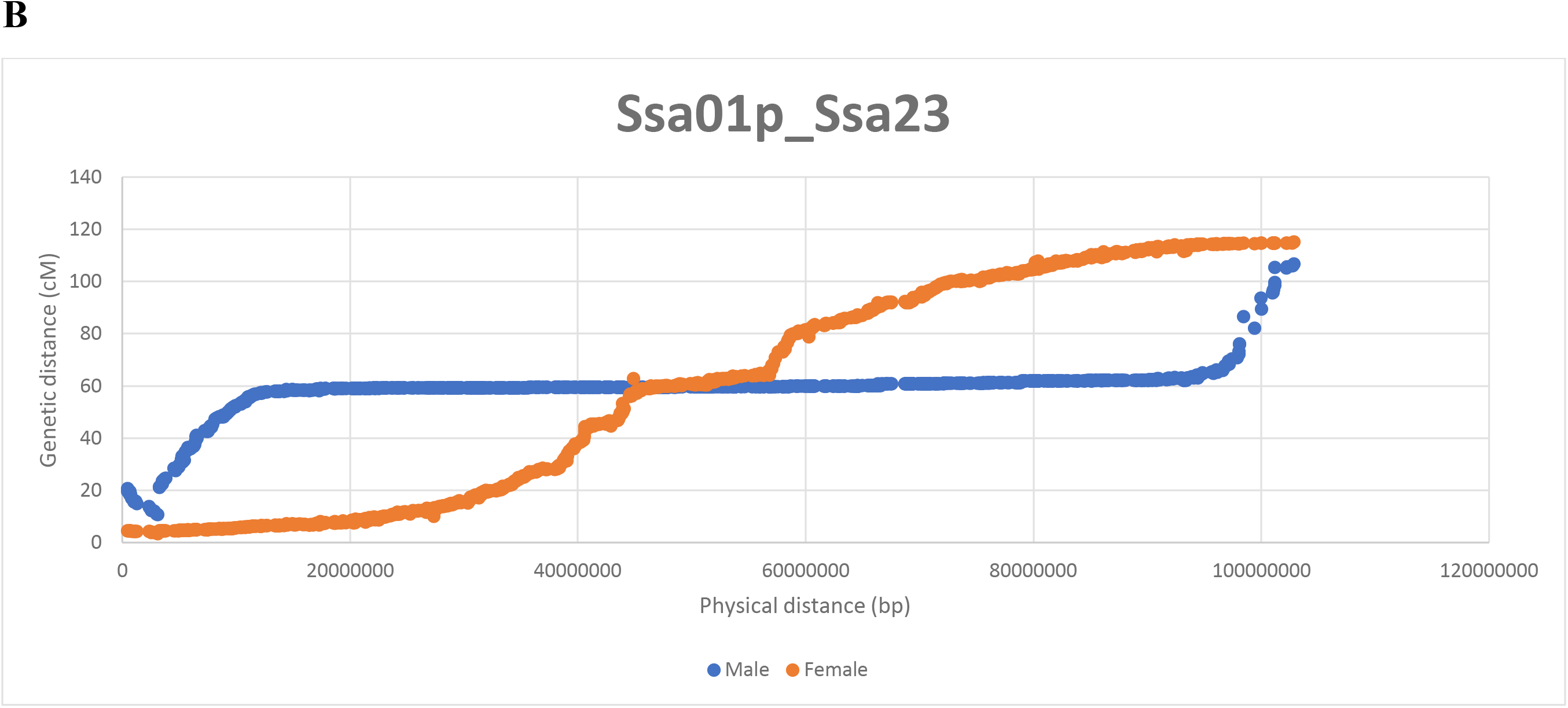
Genetic distance is plotted against physical distance to illustrate the differences in the typical recombination patterns between males and females in salmonids. In the acrocentric chromosome (A) the paternal chromosome displays almost no recombination except for the area proximal to the q-arm telomere. Conversely, the maternal pattern displays a higher rate of recombination near the centromere, a continuous even level of recombination throughout the rest of the chromosome, and gradual decrease to a plateau near the telomere. In the metacentric chromosome (B) the maternal chromosome displays strong recombination interference in the centromeric region and then similar pattern to the acrocentric chromosome in the direction of either telomere, while the paternal pattern is consistent with high interference in recombination except for elevated levels of recombination near both telomeres.

### Genome assembly

The genome assembly presented in the present research is based on DNA from a single male from the USDA St. John River stock of N. A. Atlantic salmon that was sequenced using PacBio long-read sequencing technology. The PacBio sequences were assembled into contigs with Canu and those contigs were joined into scaffolds using Bionano optical mapping and Hi-C proximity ligation sequence data. Using genetic linkage information, the scaffolds from the Hi-C assembly were then anchored to and ordered on chromosomes to generate the chromosome sequences of the USDA_NASsal_1.1 assembly (GCA_021399835.1).

The Canu sequence assembly was composed of 14,845 contigs with a total length of ~3.97 Gb, N50 of 1.45 Mb, and BUSCO score of 95.8% (using actinopterygii_odb9 as the lineage dataset). While the estimated genome size of Atlantic salmon is ~3.0 Gb (Lien et al. 2016) it was clear that heterozygosity in this individual genome caused duplication of haplotypes for some of the heterozygous genomic regions. After the purging steps removed duplicated haplotypes resulting from heterozygous sequences, the BUSCO score was improved slightly to 96.0%, but the total length of the assembly contigs at ~3.89 Gb was still greater than the expected genome size. More detailed statistics for each step in the assembly pipeline are shown in **Table 2**. The Bionano hybrid scaffolding pipeline used the optical map to join neighboring or overlapping Canu contigs and to break chimeric sequence contigs that were misassembled by Canu. The final Bionano scaffolded assembly included 2,012 scaffolds of joined primary contigs and haplotigs and primary contigs that could not be joined into scaffolds. After this step the total length of the assembly was finally reduced to within the expected genome size at ~2.83 Gb and the N50 increased to 11.9 Mb (**Table 2**). The Hi-C ligation proximity mapping further reduced the number of scaffolds to 1,633 and increased the N50 to 37.9 Mb and the BUSCO score to 96.2%, with negligible impact on the total length of the assembly (**Table 2**).

**Table 2.** Statistics of the North American Atlantic salmon genome assembly at each step of the assembly pipeline.

| **Feature** | **Canu contigs** | **Purge duplicates** | **BioNano scaffolds** | **Hi-C scaffolds** | **Chr. Sequences <sup>1</sup>** |
| --- | --- | --- | --- | --- | --- |
| Number of scaffolds/contigs | 14,845 | 14,240 | 2,012 | 1,633 | 1,755 |
| Total length (Mb) | 3,970 | 3,889 | 2,835 | 2,834 | 2,829 |
| Maximum length (Kb) | 24,768 | 24,768 | 70,926 | 115,072 | 92,895 |
| Minimum length (bp) | 1,003 | 1,012 | 3,771 | 3,768 | 3,768 |
| N50 (Kb) | 1,452 | 1,534 | 11,936 | 37,905 | 16,202 |
| BUSCO <sup>2</sup> | 95.8% | 96.0% | 95.8% | 96.2% | 96.2% |

<sup>1</sup> Includes the unplaced scaffolds (N=1,122). Total number of scaffolds mapped to chromosomes was 633. Some of the largest scaffolds were broken by the linkage map information (e.g. two parts were mapped to two different chromosomes).
<sup>2</sup> BUSCO: Benchmarking Universal Single-Copy Orthologs. Showing the percent of complete protein-coding genes detected (using actinopterygii\_odb9 as the lineage dataset).

Using information from the new linkage map to anchor the scaffolds to karyotype chromosomes broke some of the largest scaffolds due to their alignment with different linkage groups. Overall, 633 scaffolds were anchored to 27 linkage groups with 914 spanned gaps and 606 un-spanned gaps to generate 27 chromosome sequences in a total length of ~2.53 Gb, or 89% of the total genome assembly length. The distribution of chromosomes by physical length in base-pairs from this assembly is shown in comparison with the chromosomes of the European Atlantic salmon reference genome assembly in **Figure 4**. The total length of the 1,122 unplaced scaffolds that are not anchored to a chromosome is ~300 Mb. The total number of scaffolds in the chromosome level assembly for the N.A. Atlantic salmon is therefore 1,755, total length is ~2.83 Gb, total un-gapped sequence length is ~2.79 Gb, and scaffold N50 and L50 are 16.2 Mb and 42, respectively.

**Figure 4.**
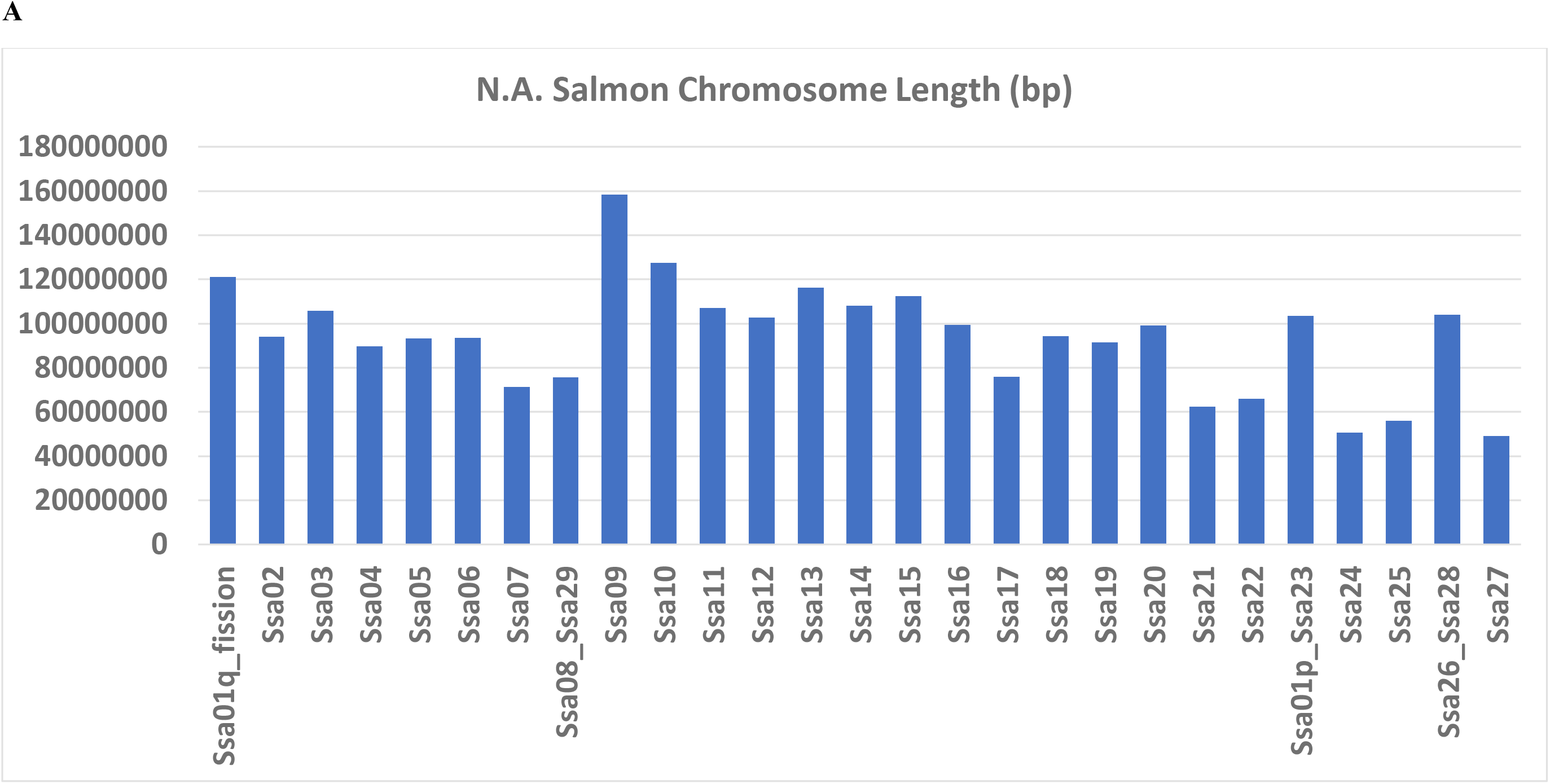
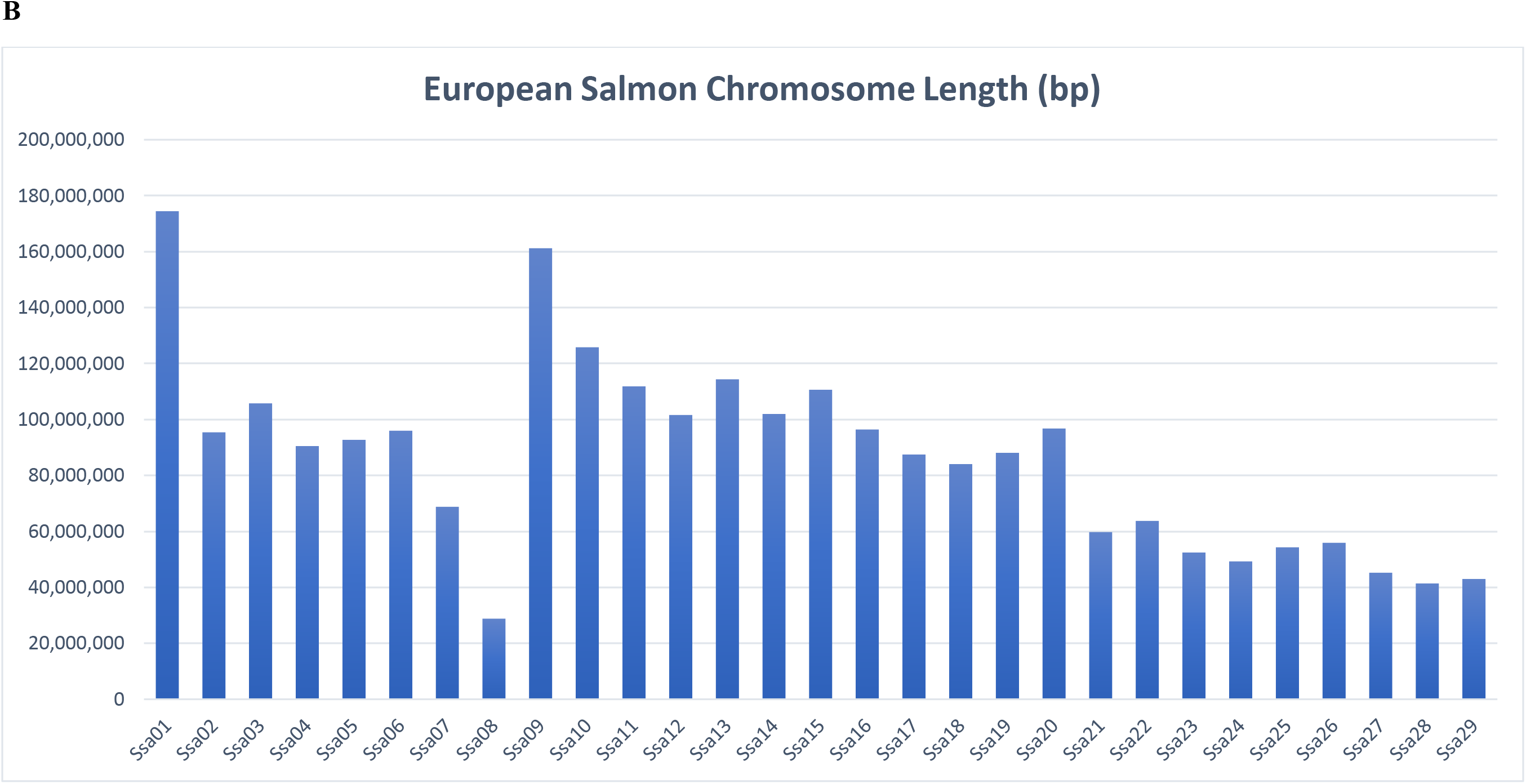
Distribution of chromosome lengths in base-pairs in the assembly from N.A. Atlantic salmon (accession GCA_021399835.1) presented in this report (A) vs. (B), the distribution in the reference genome assembly from the European lineage (Ssal_v3.1; GCA_905237065.2).

A comparison of the assembly statistics between this assembly and the two annotated reference genome assemblies for the European Atlantic salmon lineage is shown in **Table S4**. The contiguity of this assembly is much better than the previous assembly (GCA_000233375.4) {Lien, 2016 #4769} which had scaffold N50 and L50 of ~0.6 Mb and 594, respectively. However, the contiguity of the new reference for the European Atlantic salmon lineage genome assembly (GCA_905237065.2) with scaffold N50 and L50 of 28 Mb and 33, respectively, is better than the assembly we present here. It also has a nearly perfect BUSCO score showing 99.3% of complete protein-coding genes detected compared to 96.2% in this genome assembly for the N.A. Atlantic salmon lineage.

### SNP array positions on the chromosome sequences

The genome positions of the SNPs from the Axiom 50K array that were mapped to unique positions of the N.A. Atlantic salmon chromosome sequences with no indels and including 60 bp flanking sequences for each SNP are available in **Supplemental File S3**. The distribution of the 50K SNPs on the chromosomes of the genome assembly is shown in **Figure S1**. The average number of SNP markers per chromosome is 1,724.7 with a minimum of 920 and a maximum of 2,885. An additional 188 SNPs were mapped to unplaced scaffolds or contigs. The SNP marker density per 1 Mb in each of the 27 chromosomes is diagrammed in **Figure 5**. The marker density range is from zero to 40 per 1 Mb. The marker density fluctuates within the chromosomes with lower density typically near the centromeres and telomeres.

**Figure 5.**
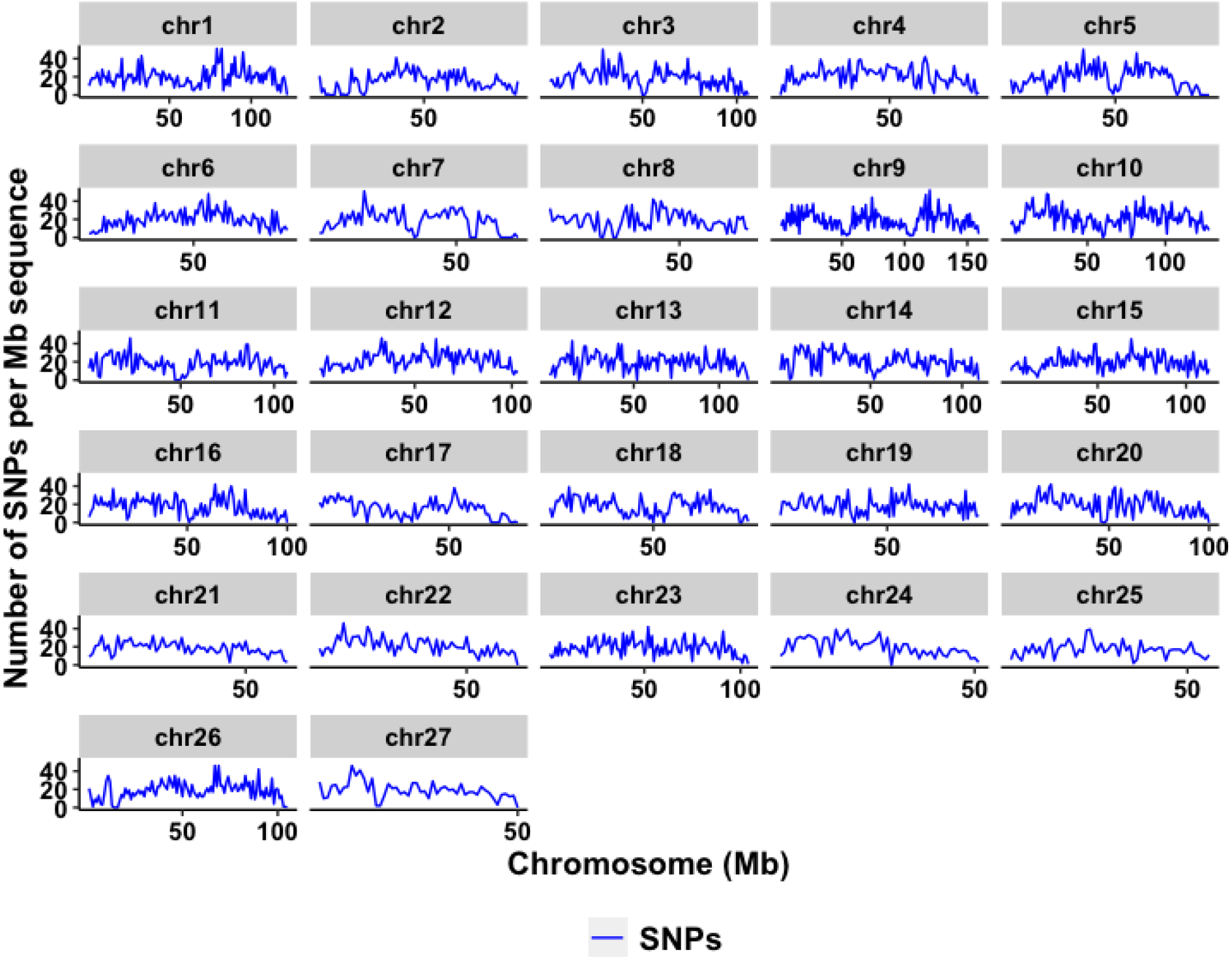
Density of markers from the N.A. Atlantic salmon Axiom 50K SNP array per million base pairs throughout the genome assembly chromosomes.

### Comparison between European and N.A. genome assemblies

Similar to rainbow trout’s plasticity in chromosome number between populations which correlates with geographic distribution (Thorgaard 1983; Gao et al. 2021), the Atlantic salmon genome also exhibits tolerance to chromosomal rearrangements. While other karyotypes have been reported for N.A. Atlantic salmon from other populations with chromosome numbers varying from 2N=54 to 2N=58 (see discussion in Brenna-Hansen et al., 2012), here we found unequivocal evidence that supports 2N=54 in the SJR strain individual fish that we used for the genome assembly presented in this report. The results of our linkage analysis coupled with the de-novo assembly of large sequence scaffolds confirmed the chromosomal rearrangements between N.A. and European Atlantic salmon proposed previously by Brenna-Hansen et al. based on their own linkage analysis and karyotyping (Brenna-Hansen et al. 2012).

The chromosome sequences from this assembly of the N.A. salmon genome (USDA_NASsal_1.1; GCA_021399835.1, N=27) were aligned with the chromosome sequences of the reference genome from the European lineage (Ssal_v3.1; GCA_905237065.2, N=29) to identify and confirm the chromosomal rearrangements. Graphic representations of the alignments for all 27 chromosomes can be found in Supplemental **File S4**. For 23 of the 27 chromosomes, we found simple collinear sequence alignments indicating highly similar order and direction of those chromosome sequences and the genes that they carry as illustrated in **Figure S2**. The rearrangements we found in the remaining four chromosomes can be explained by four major separate events. The first was fission in the centromere region of chromosome Ssa01 resulting in the separation of chromosome arms Ssa01q and Ssa01p. The second was the translocation and fusion of Ssa01p with Ssa23 resulting in Chr 23 of the N.A. lineage genome assembly. The evidence from sequence alignments for those two chromosomal rearrangements is illustrated in **Figure 6**. According to Brenna-Hansen et al. (2012) this is the most common rearrangement between salmon of European and N.A. origins as all the chromosome karyotypes that they have observed from N.A. salmon were homozygous for the translocation of Ssa01p to Chr 23. The third rearrangement event is the formation of Chr 08 in the N.A. genome from the fusion of the acrocentric chromosomes Ssa08 and Ssa29 (**Figure 7**). The fourth is the fusion of Ssa26 and Ssa28 to form Chr 26 in the N.A. genome assembly (**Figure 8**). According to Brenna-Hanson et al. (2012), their karyotype preparations revealed heterozygosity to the latter two chromosomal fusions (Ssa08/29 and Ssa26/28) in different individuals of N.A. origin, indicating that those karyotypes may not be shared by all populations of Atlantic salmon from the N.A. lineage. Hence, it is likely that those two chromosomal fusion events are more recent than the translocation event of Ssa01p to Chr 23 in N.A. Atlantic salmon.

**Figure 6.**
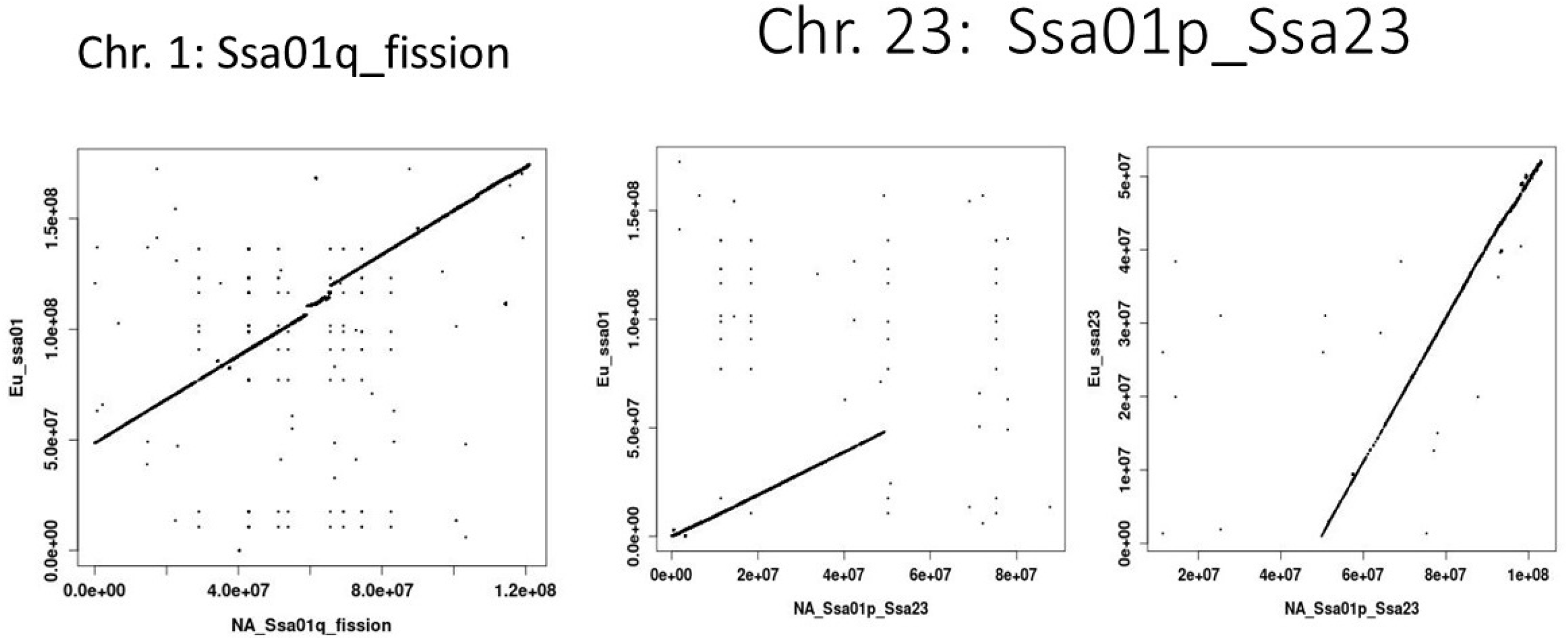
Chromosome fusions and fissions between the European and N.A. Atlantic salmon lineages. Fission of the European chromosome Ssa01q and p arms and fusion of Ssa01p with Ssa23 in the N.A. Atlantic salmon genome. Sequences were compared using data from the reference assembly for the European lineage in NCBI (Ssal_v3.1; GCA_905237065.2).

**Figure 7.**
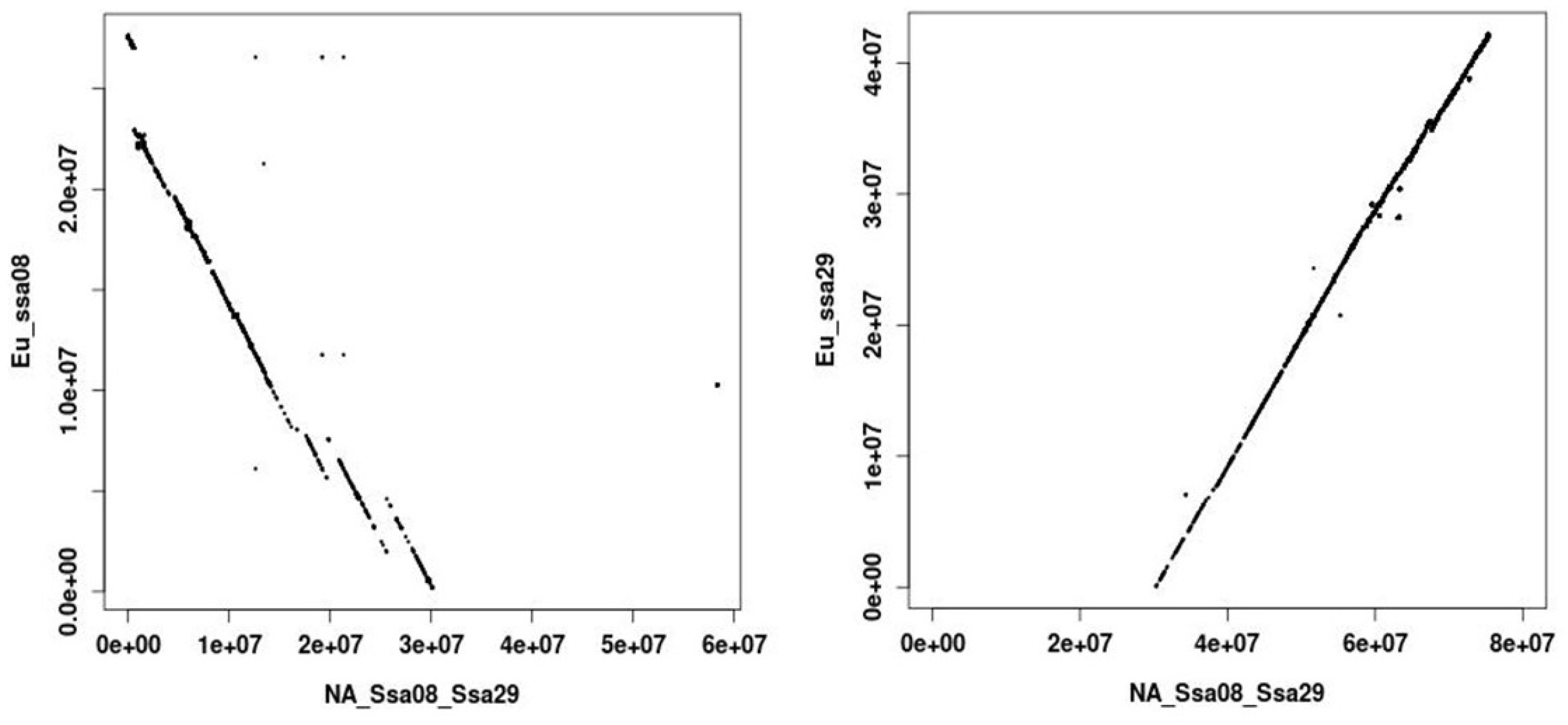
Fusion of the European lineage chromosomes Ssa08 and Ssa29. Sequences were compared using data from the reference assembly for the European lineage in NCBI (Ssal_v3.1; GCA_905237065.2).

**Figure 8.**
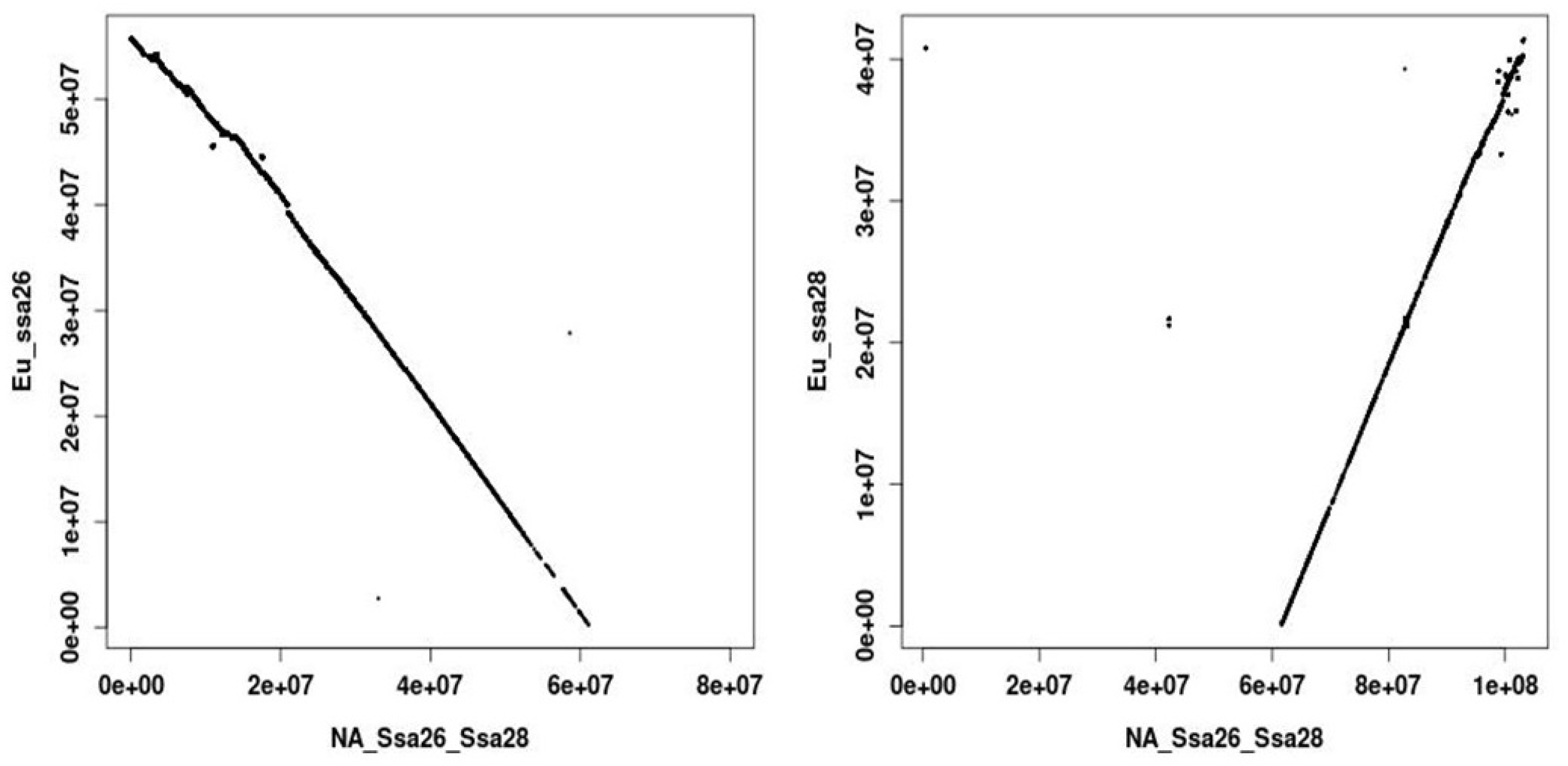
Fusion of the European lineage chromosomes Ssa26 and Ssa28. Sequences were compared using data from the reference assembly for the European lineage in NCBI (Ssal_v3.1; GCA_905237065.2).

### Annotation via alignment with European lineage reference genome in NCBI

Gene annotation for this genome assembly accession is not available currently from NCBI or Ensembl. Chromosome sequences or specific regions of interests from the genome assembly can be aligned by Blast searches with the Refseq annotated version of the European lineage genome assembly (GCF_905237065.1). For example, the T cell receptor beta 01 region which is found on Ssa01p in the European lineage reference genome has been shown to be translocated to Chr 23 (Ssa01p_Ssa23) in the assembly of the N.A. Atlantic salmon genome presented here (Grimholt et al. 2022). In addition, the alignments of USDA_NASsal_1.1 (GCA_021399835.1) to the annotated Ssal_v3.1 (GCF_905237065.1) genome assembly are now available in the NCBI remap menu at: https://www.ncbi.nlm.nih.gov/genome/tools/remap. The USDA assembly can be used as the source assembly and the Ssal_v3.1 as the target assembly. After pasting or uploading the chromosome coordinates of interest from the source assembly the user can choose the output format of interest. The output file in genome workbench format can be used to view the aligned chromosome region or regions of choice in a genome browser.

### Conclusions

We report here the de-novo assembly of the first chromosome level reference genome and the development and validation of the first publicly available high density SNP array for the North American lineage of Atlantic salmon. These genomic resources are required for implementing genomic selection for aquaculture production and are crucial to current research on the genetics, biology, evolution, and ecology of this sub-species that holds high economic and social value in Eastern United States, Canada and Australia.

## Acknowledgements

We thank Kristy Shewbridge for her technical assistance with the preparation of the DNA sample used for genome sequencing. We thank Dr. Serap Gonen and Dr. Matthew Baranski from Mowi for sharing SNP polymorphism information from the European SNP array and for contributing samples of farmed salmon from European origin. We thank Dr. Matthew Kent for sharing the flanking sequence data of the markers used in the linkage map published by Brenna-Hansen et al. (2012). This study was supported by the USDA Agricultural Research Service in-house project numbers 8030-31000-004/005 and 8082-31000-012/013 and 6066-31000-016. Mention of trade names or commercial products in this publication is solely for the purpose of providing specific information and does not imply recommendation or endorsement by the U.S. Department of Agriculture. USDA is an equal opportunity provider and employer.

## Authors’ Contributions

Guangtu Gao: Writing – original draft, Conceptualization, Methodology, Formal analysis, Data curation, Writing – review & editing. Geoffrey C. Waldbieser: Investigation, Methodology, Resources, Writing – review & editing. Ramey C. Youngblood: Investigation, Methodology, Formal analysis. Dongyan Zhao: Writing – original draft, Methodology, Formal analysis. Michael R. Pietrak: Project administration, Investigation, Validation. Melissa S. Allen: Validation, Data curation. Jason A. Stannard: Project administration, Methodology, Validation. John T. Buchanan: Methodology, Supervision. Roseanna L. Long: Investigation, Validation. Melissa Milligan: Investigation, Validation. Gary Burr: Investigation, Validation. Moira J. Sheehan: Resources, Supervision. Brian E. Scheffler: Resources, Supervision. Caird E. Rexroad III: Conceptualization, Resources. Brian C. Peterson: Conceptualization, Supervision, Resources. Yniv Palti: Writing – original draft, Conceptualization, Methodology, Supervision, Project administration, Resources, Writing – review & editing. All authors read and approved the manuscript.

## Data availability statement

The reference genome is available for downloading and browsing at NCBI (GenBank: GCA_021399835.1; BioProject: PRJNA759632). The PacBio long reads and the Illumina short reads raw sequence data used for the genome assembly are in NCBI-SRA (under the same BioProject; SRR accession awaited processing). The new SNP discovery dataset, the linkage map and the genome positions of the SNPs from the 50K Axiom array are available as supplemental files. All other datasets used and/or analyzed in the current study are available from the corresponding author on reasonable request.

---

**Table S1.** Populations samples included in the validation of continent of origin markers. In Excel file.

**Table S2.** Markers validated for identification of the Atlantic salmon fish continent of origin. In Excel file.

**Table S3.** Description and flanking DNA sequence information for the SNP markers used for sex determination in the 50K SNP array. In Excel file.

**Table S4:** Comparison of genome assembly statistics with the two reference version assemblies from the European lineage of Atlantic salmon.

| **Feature** | **European Old<sup>1</sup>** | **European New<sup>2</sup>** | **North American<sup>3</sup>** |
| --- | --- | --- | --- |
| Number of scaffolds | 839,389 | 4,222 | 1,755 |
| Total length (Mb) | 3,412 | 2,756 | 2,829 |
| Scaffold N50 (Kb) | 592 | 28,058 | 16,202 |
| Scaffold L50 | 594 | 33 | 42 |
| Contig N50 (Kb) | 36 | N/A | 4,207 |
| Contig L50 | 14,749 | N/A | 150 |
| No. of Chromosomes | 29 | 29 | 27 |
| BUSCO <sup>4</sup> | 97.9% | 99.3% | 96.2% |

<sup>1</sup> NCBI Genbank accession GCA\_000233375.4.
<sup>2</sup> NCBI Genbank accession GCA\_905237065.2.
<sup>3</sup>NCBI Genbank accession GCA\_021399835.1.
<sup>4</sup> BUSCO: Benchmarking Universal Single-Copy Orthologs. Showing the percent of complete protein-coding genes detected (using actinopterygii\_odb9 or \_odb10 as the lineage dataset).

**Figure S1.**
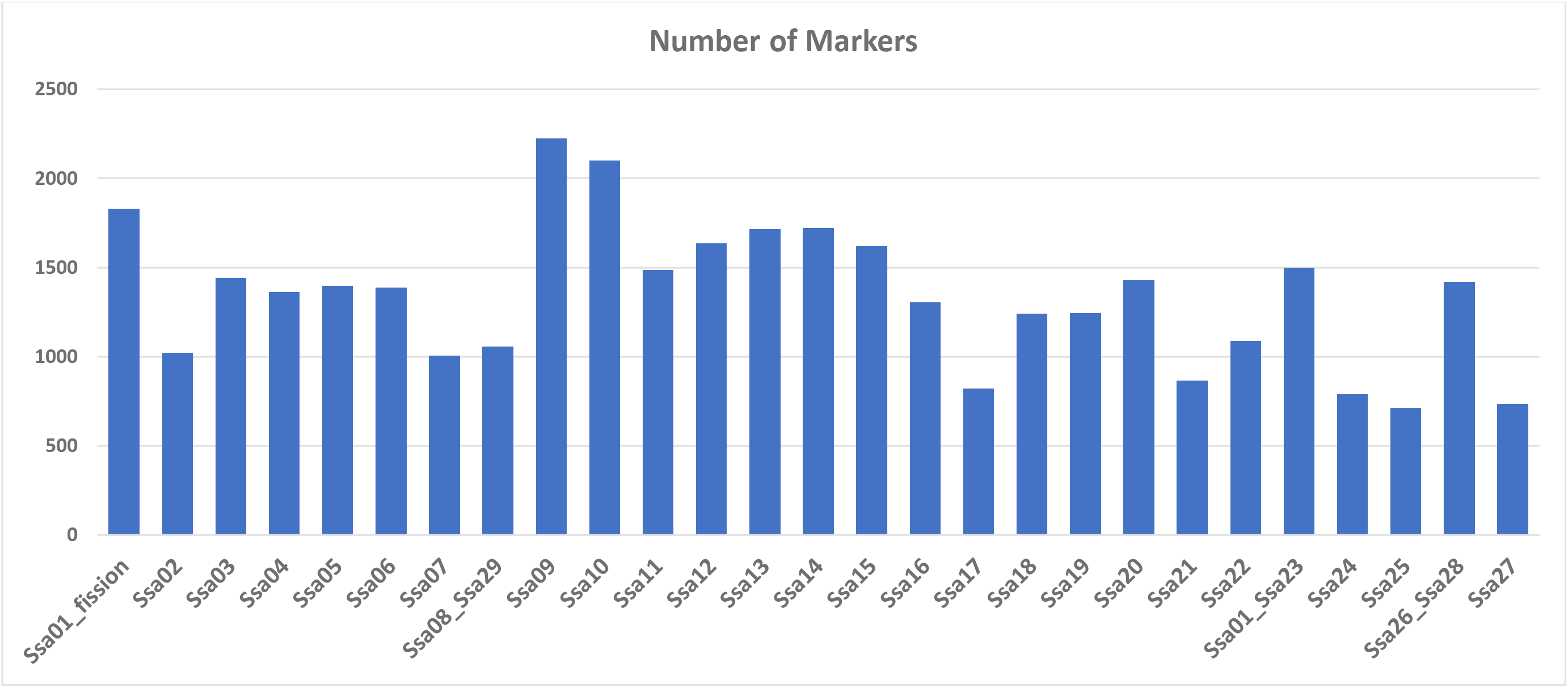
Distribution of markers from the N.A. Atlantic salmon Axiom 50K SNP array on the genome assembly chromosomes.

**Figure S2.**
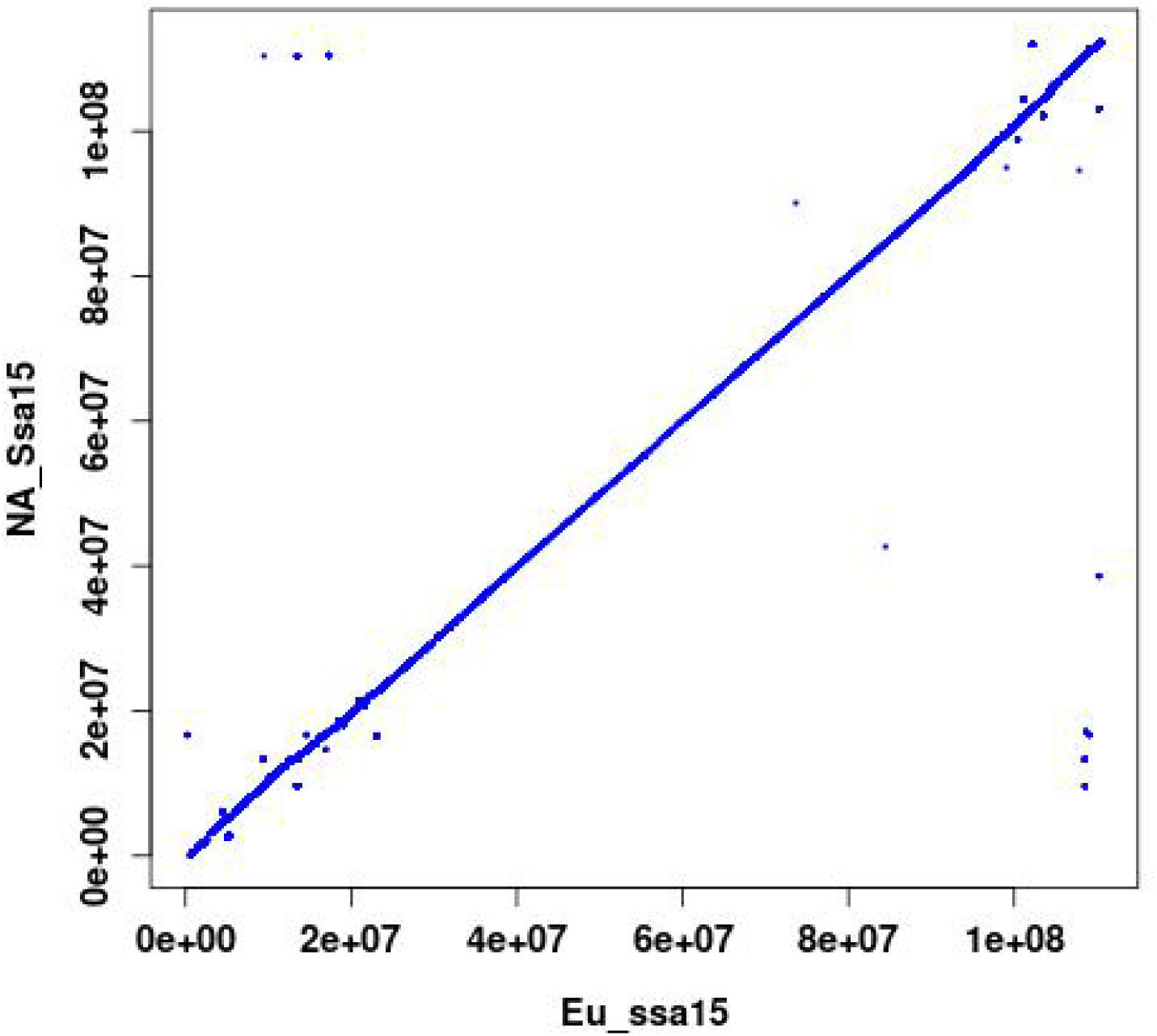
A typical linear alignment between chromosome sequences from the N.A. and European Atlantic salmon genomes is shown for chromosome Ssa15. Sequences were compared using data from the reference assembly for the European lineage in NCBI (Ssal_v3.1; GCA_905237065.2).

## Supplemental Files

**Supplemental File 1:** New database of ~2.3M uniquely mapped SNPs generated for N.A. Atlantic salmon including chromosome positions and alternative alleles. (Available to download from GSA journals Figshare and from Github: https://github.com/guangtugao/Na_salmon_data/blob/main/File%20S1%20-%20DB%20SNPs%20Unique_U.csv.zip)

**Supplemental File 2:** Linkage map of north American Atlantic salmon from the St. John River aquaculture strain.

**Supplemental File 3:** Genome positions of the 50K SNPs included in Axiom high-density SNP array.

**Supplemental File 4:** DNA sequence alignments between the Atlantic salmon chromosomes from the north American lineage genome assembly (this report; accession GCA_021399835.1) and the European lineage reference genome assembly (Ssal_v3.1; GCA_905237065.2).

